# Limitations of Genomic Foundation Models for Decoding Regulatory Mechanisms in ALS

**DOI:** 10.64898/2026.10.01.756029

**Authors:** Arghamitra Talukder, Alan Kaplan

## Abstract

Genomic foundation models promise to infer regulatory consequences of genetic variants directly from DNA sequence, but how well these predictions translate to clinical utility remains unclear. To probe this question, we evaluate AlphaGenome (AG) for understanding amyotrophic lateral sclerosis (ALS) genetics without any disease-specific fine-tuning across three settings of increasing difficulty: recovering measured variant effects, reconstructing a known splicing mechanism, and generating de novo regulatory hypotheses at unresolved loci. At population scale, AG failed to recover measured effect sizes across 1,254 spinal-cord splicing quantitative trait loci (QTL) (Spearman = 0.041, Pearson = 0.104) and was precise only at negligible recall (precision = 1.00 at 1.3% recall). At a mechanistically resolved locus of *UNC13A* cryptic-exon event, AG predicted the intronic risk variant’s splicesite usage opposite its known risk effect (Δ ≈ −2 × 10^−6^), showed near-identical effects in disease-relevant and disease-irrelevant tissues (cerebellar effects reached ~60% of neural magnitude), and recovered only a partial cis-regulatory signature. Finally, AG showed limited gene-level resolution across seven ALS genome-wide association study (GWAS) loci, with predicted effects declining sharply as a gene lay farther from the variant (a 10-fold increase in distance cost 0.95 standard deviations of signal; *p* = 2.7 × 10^−23^). In summary, AG can identify a small set of high-confidence regulatory variants, but cannot reliably rank variants by biological effect, or assign regulatory signals to the correct effector gene at unresolved loci. Thus, although AG captures meaningful local regulatory effects, these predictions do not resolve the effect sizes and effector genes needed to connect ALS-associated variants to their causal mechanisms.

## 1 Introduction

Biology is entering an era of foundation models [19, 10, 11, 29], increasingly envisioned as unified, general-purpose systems that could reduce reliance on task-specific models and pipelines [2, 22, 30], costly experimentation [7, 12], and in some cases on expert analysis [3]. A growing class of genomic foundation models [1, 5] extends this ambition to gene regulation, predicting molecular function from DNA sequence at single-base-pair resolution while integrating context across up to a megabase. A single model spans diverse regulatory readouts, transcription, splicing, chromatin accessibility, transcription-factor binding, chromatin state, three-dimensional genome organization, and scores how single-nucleotide variants perturb them [14]. Because these modalities are governed by shared regulatory logic linked through a causal cascade, jointly predicting them is often taken as evidence of the model acquiring a genuine mechanistic understanding of biology, rather than merely sophisticated memorization of its training data.

Disease-associated regulatory variation offers a particularly stringent test of this premise, requiring models to connect sequence perturbations to the genes, cellular contexts, and regulatory mechanisms through which they influence pathology. At the same time, this is precisely where genomic foundation models could have their greatest scientific and clinical value, provided their predictions are mechanistically faithful. Amyotrophic lateral sclerosis (ALS) provides such a setting. ALS a fatal neurodegenerative disease affecting one in 350 individuals is characterized by progressive degeneration of upper and lower motor neurons, leading to muscle weakness, paralysis, and ultimately respiratory failure [15, 28]. In some cases, ALS is driven by rare high-effect variants. Sporadic ALS instead behaves as a complex trait, with an estimated heritability of 40–50% [24, 26]. This complexity poses several challenges for understanding how genetic variation contributes to disease. The first challenge is *causal variant identification*. Pinpointing causal ALS variants is hard because risk is driven not by isolated variants but by many interacting partners acting through cell-type-specific noncoding regulatory regions, where the bulk of heritability lies yet variant-by-variant genome-wide association study (GWAS) remain systematically underpowered [31, 9, 27]. Despite substantial effort to fine-map ALS risk loci and prioritize candidate variants [33, 28], a second challenge remains: *mechanistic resolution*. Noncoding regulatory elements remain sparsely annotated, so linking a variant to its target gene and cellular context has required painstaking, locus-by-locus experimental work [18, 32].

In this work, we ask whether a genomic foundation model can close this gap. We evaluate AlphaGenome (AG) [1] zero-shot—without any ALS-specific training or task-specific fine-tuning—to determine whether its learned predictions of noncoding variant effects on molecular regulation can yield novel biological insight into ALS. Together, these analyses reveal how far a general-purpose genomic foundation model can support disease interpretation, defining both the confidence its predictions warrant and the specificity with which they resolve variant effects. We conduct this assessment in three parts:

- **Quantitative fidelity**. We assess whether AG can quantitatively recover the experimentally measured splicing effects of 1,254 ALS spinal-cord splicing quantitative trait locus (sQTLs). Accurate effect-size prediction is a prerequisite for *causal variant identification*, particularly for rare variants beyond the reach of population-based measurement. Instead, predicted magnitude is largely unrelated to measured effect (Spearman = 0.041, Pearson = 0.104), yielding reliable signal for only a small subset (precision reaches 1.00 only at 1.3% recall) and undermining its use to prioritize unmeasured, particularly rare, variants by regulatory impact.
- **Mechanistic fidelity**. We evaluate whether AG can reconstruct a known disease mechanism using the *UNC13A* cryptic-splicing. This benchmark provides a direct validation of whether AG predicts sufficiently *reliable mechanistic interpretation* of disease-associated variants. AG recovers a partial cis-regulatory signature (the risk variants raise cryptic-site splice-site usage to 7.6 × 10^−5^ versus *<* 3 × 10^−8^ for the neutral variants); it fails to reproduce variant direction (the intronic risk variant yields opposite effect ≈ −2 × 10^−6^), and tissue specificity.
- **De novo inference**. We finally ask whether AG can generate specific regulatory hypotheses at ALS GWAS loci where the mechanism remains unresolved. This tests the model’s utility for *mechanistic resolution*—generating novel insight into disease mechanism where no established ground truth exists. Instead, gene-level signal is largely explained by proximity to the lead variant (active positions span only ~5–28 kb of the supplied 1-Mb context), limiting reliable effector-gene prioritization.

## 2 Result

In following sections we evaluate the PyTorch implementation of AlphaGenome [4] along the three axes introduced above. We focus on the limits and failure modes that emerge as biological interpretation becomes increasingly demanding. Methods and data are detailed in Appendix B.

### 2.1 Quantitative fidelity: Population-scale evaluation on ALS sQTLs

We benchmarked AG’s predictions (Eq. 1) against sQTLs in cervical spinal cord from the NYGC ALS Consortium [16] to assess whether it can recover experimentally observed variant effects (*β*).

To test whether AG distinguishes regulatory variants from comparable non-regulatory ones, each true sQTL defined as the lead single-nucleotide polymorphism (SNP) per junction *q <* 0.01 was paired with five non-sQTL variants drawn from the same junction and matched on variant–junction distance and allele frequency, excluding variants in linkage disequilibrium (LD) with the sQTL (Appendix B.1.2, Fig. 5a). We retained the 1,254 junction groups whose variant and both splice sites fell within 131,072,bp input window and had a valid AG’s splice-junction output. Details are provided in Appendix B.1.

We first asked whether AG predicts the magnitude and direction of measured sQTL effects. Relating the normalized splice-junction score (Norm. SJ Δ) explained in Eq. 3 to the measured sQTL effect size (*β*) reveals no meaningful correspondence (Fig. 1a; Spearman = 0.041, Pearson = 0.104). Variants with near-zero predicted effects (71.3% with |Norm. SJ Δ| *<* 1 × 10^−4^) span nearly the full range of measured effects (|*β*| up to 2.54), while the few variants with large predicted effects cluster at modest positive *β*. One explanation is that AG assigns large effects based on the variant’s proximity to a splice site. Consistent with this, |Norm. SJ Δ| decreased with the variant’s distance to the nearest splice site (Fig. 1b); Spearman = −0.30), with the median predicted effect dropping about 60-fold from the nearest to the farthest distance bin.

**Figure 1.**
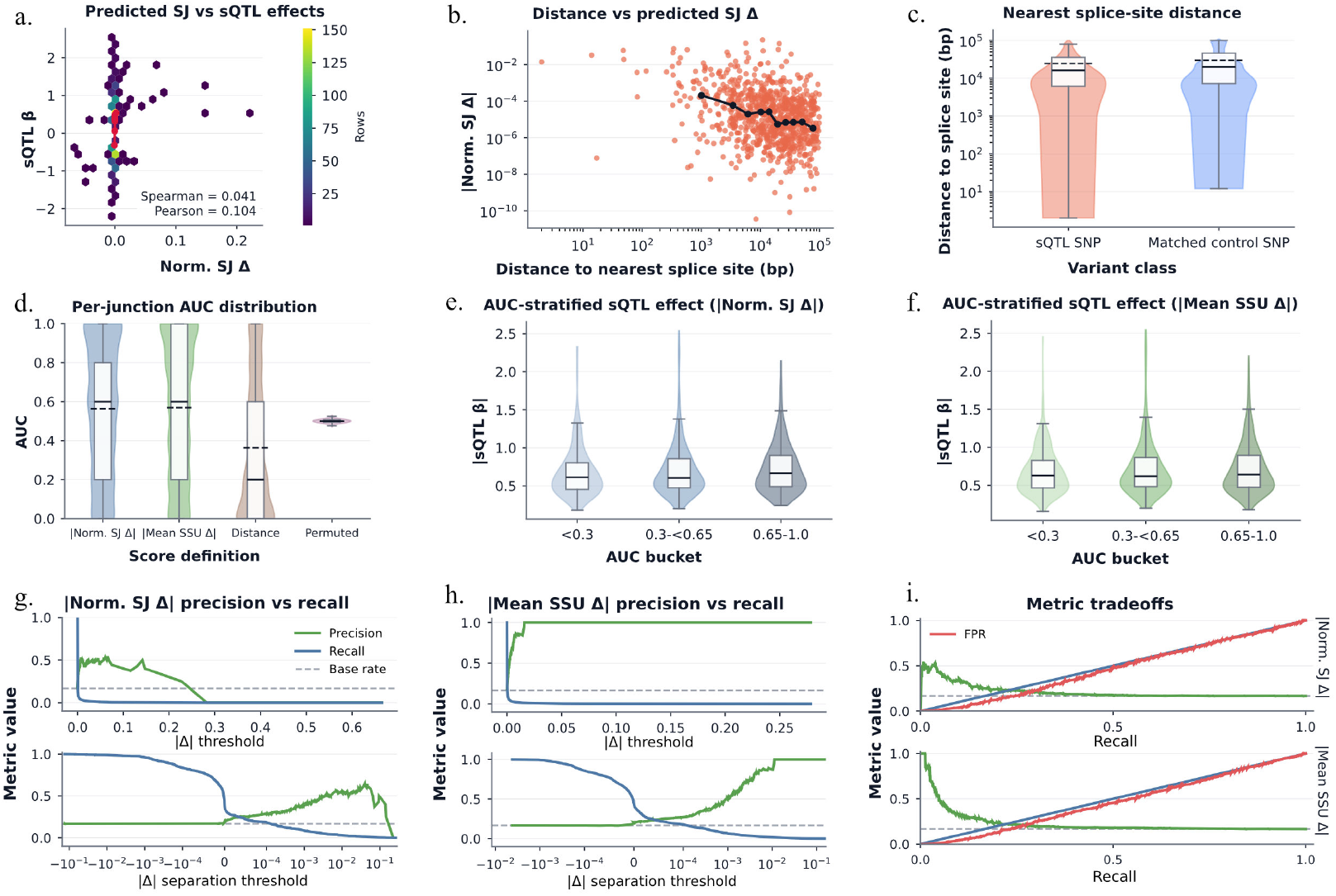
Benchmarking AG on ALS sQTLs. AG is benchmarked against ALS cervical spinal-cord sQTLs for regulatory variant-effect prediction. **(a)** Relationship between the normalized AG-predicted splice-junction effect (Norm. SJ Δ) and the measured sQTL effect size (*β*), shown as a hexbin density. **(b)** Predicted splice-junction effect magnitude as a function of variant distance to the nearest splice site. **(c)** Distance distribution to the nearest annotated splice site for true sQTL versus matched control variants. **(d)** Distribution of per-junction discrimination (AUC) for distinguishing sQTL variants from matched controls using normalized splice-junction change (|Norm. SJ Δ|), mean splice-site-usage change (|Mean SSU Δ|), a distance heuristic, and permuted labels. Within each box the solid line marks the median and the dashed line the mean. **(e, f)** Distribution of measured |*β*| stratified by per-junction AUC bucket for |Norm. SJ Δ| (e) and |Mean SSU Δ| (f). **(g, h)** Precision and recall as a function of |Δ_AG_| threshold (top) and |Δ|_sep_ threshold (bottom) for |Norm. SJ Δ| (g) and |Mean SSU Δ| (h); dashed line is the base rate. **(i)** Recall-indexed trade-offs among classification metrics for |Norm. SJ Δ| (top) and |Mean SSU Δ| (bottom).

We next asked a less stringent question. Even if the predicted effect does not quantitatively reproduce the measured sQTL effect, does AG assign a larger perturbation to the experimentally supported sQTL (case/positive variants) than to matched non-sQTL (control/negative variants)? Because splice-site proximity could confound this comparison, we first verified that the matched controls were not systematically farther from the junction than the true sQTLs. We verified this at the distribution level. The two classes showed broadly overlapping distance distributions, with true sQTLs only marginally closer than controls (median 16.1 vs. 19.7 kb; Fig. 1c). With this proximity balance established, we evaluated a within-junction ranking using the absolute prediction magnitude, |Norm. SJ Δ| (Eq. 3) and |Mean SSU Δ| (Eq. 4), and calculated an AUC independently for each junction group. Both AG scores modestly exceeded chance (median AUC 0.60), while distance to the nearest splice site performed below chance (median 0.20) and label permutation remained expected narrow distribution around 0.5 (Fig. 1d). The scores thus carry genuine signal. However, performance varied substantially across junctions (interquartile range 0.20–0.80). Thus, the modest aggregate enrichment does not reflect uniformly successful identification of sQTLs. We further asked whether this heterogeneity could be explained by *β*. We therefore stratified junctions by the AUC and examined the corresponding experimental effect sizes. Although a small shift toward larger effects is visible in the higher-AUC groups (median |*β*| rises only from 0.61 to 0.67 across AUC buckets for |Norm. SJ Δ| and from 0.63 to 0.64 for |Mean SSU Δ|), there is no strong separation (Fig. 1e,f). So junctions the model ranks well do not carry systematically larger true effects. Consistently, measured |*β*| remained largely unchanged across predicted-separation strata, with within-stratum correlations remaining near zero (Appendix Fig. 5b).

The broad AUC distributions suggested that AG might still be useful when restricted to variants receiving particularly strong or well-separated predictions. We therefore examined precision and recall as the decision threshold was varied in Fig. 1g,h, both by predicted effect magnitude (|Δ_AG_|) and by signed separation (|Δ|_sep_) defined in Eq. 1 and 2. For both readouts, precision rises above the base rate (0.17) only at the most stringent |Δ_AG_| thresholds, and exactly there recall has already collapsed toward zero: Mean SSU Δ reaches a precision of 1.00 but at a recall of only 0.013 (*n* = 16 variants), and Norm. SJ Δ peaks at a precision of 0.55 at a recall of 0.007 (*n* = 11). AG can be made precise, but only by discarding almost all true sQTLs. The |Δ|_sep_ (Fig. 1g,h, lower panels) shows the same pattern: precision increases primarily when |Δ|_sep_ is positive, meaning the predicted effect is larger for the true sQTL than for its matched controls. The two readouts diverge at the stringent end, where Mean SSU Δ precision climbs toward one while Norm. SJ Δ does not sustain it, peaking near 0.65 (*n* = 17) before collapsing to zero at the single-variant endpoint. Across the full recall range, the ROC curve follows the diagonal (AUC 0.53 and 0.54 for the two scores), indicating near-chance discrimination (Fig. 1i). AG can therefore flag a limited high-confidence subset, particularly from splice-site-usage predictions but limited to very low recall.

Across these analysis, the same limitation is evident. AG is specific but not sensitive, sufficient to nominate a variant within the extreme tail of predictions, but close to chance in ranking the majority of experimentally observed sQTLs. Moreover, the variants it identifies with high confidence are not necessarily those of greatest effect. At population scale, AG cannot prioritize the variants most likely to matter, only confirm, at low yield, that a minority are genuine.

### 2.2 Mechanistic fidelity: Evaluation at *UNC13A* cryptic splicing

The population-scale benchmark tests recovery in aggregate but not the actual mechanism. We therefore next evaluated AG’s mechanistic fidelity at a disease-associated locus where the relevant cryptic-splicing event has been experimentally characterized. We used the *UNC13A* cryptic-exon event [20, 6], in which an aberrant exon between exons 20 and 21 introduces competing cryptic splice sites. Two ALS associated variants occur in this region: rs12973192 within the cryptic exon (ES) and rs12608932 downstream of the cryptic donor (IS) (Fig. 2a). Details can be found in Appendix B.2.

**Figure 2.**
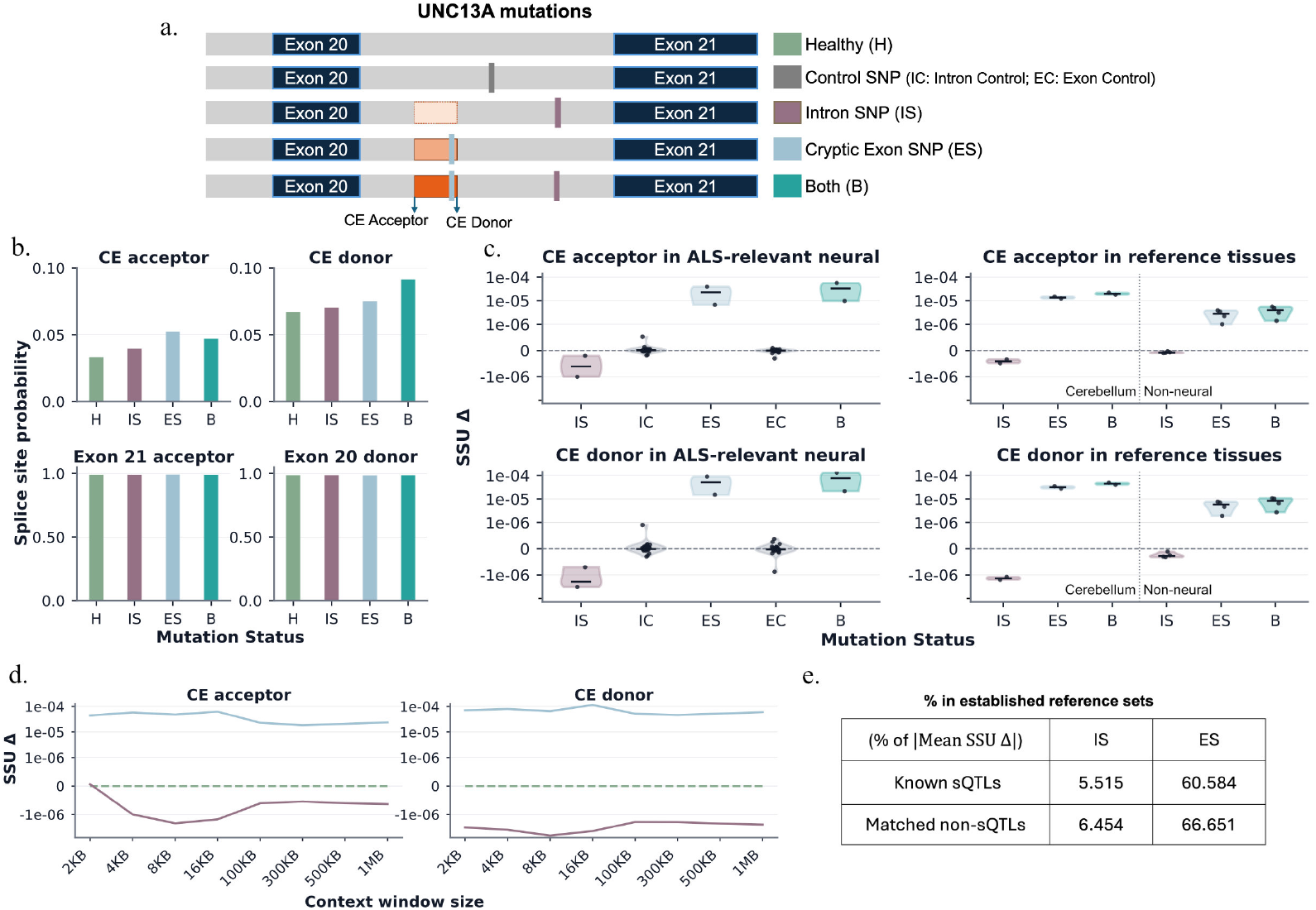
AG predictions at the *UNC13A* cryptic exon across risk-variant conditions. **(a)** Schematic of the *UNC13A* locus and the conditions scored with AG: the healthy reference (H), the intronic and cryptic-exon risk variants (IS, rs12608932; ES, rs12973192), their matched controls (IC, EC), and both risk alleles combined (B). The cryptic-exon (CE) acceptor and donor sites are indicated. **(b)** AG splice-site probabilities at the cryptic-exon acceptor and donor and the canonical exon 21 acceptor and exon 20 donor across mutation conditions. **(c)** Predicted splice site usage effect SSU Δ at the cryptic-exon acceptor (top) and donor (bottom) across ALS-relevant neural tissues (left) and non-neural reference tissues (right) across variant conditions. The violin shows the distribution across tracks. **(d)** Predicted SSU Δ at the cryptic-exon acceptor and donor across AG sequence-context window sizes, illustrating the dependence of the predicted variant effect on genomic context. **(e)** Percentile of the AG-predicted *UNC13A* IS and ES effects (% of |Mean SSU Δ|) within the distribution of |Δ_AG_| for known sQTLs and matched non-sQTLs in 2.1.

We first asked whether AG recognized the relevant local splice architecture at the *UNC13A* locus. Across variant conditions, AG assigns rising splice-site probability to the cryptic acceptor and donor as the intronic and cryptic-exon variants (IS, ES) are added (Fig. 2b). The intron variant (IS) alone produces only a small increase over the healthy baseline (H) (cryptic acceptor 0.033 →0.040, cryptic donor 0.067 →0.070). Introducing the cryptic-exon variant (ES) raises the predicted probability further (to 0.052 and 0.075), and introducing both risk alleles together (B) increases the cryptic-donor probability the most (0.091). However, the cryptic-site probabilities remain low in absolute terms (*<* 0.1) even when both variants are present, whereas the canonical exon 20 donor and exon 21 acceptor stay saturated near one across all conditions. AG therefore registers the cryptic exon as a weak alternative that never meaningfully competes with the canonical junction.

We next examined predicted splice-site usage effects (SSU Δ) in the ALS-relevant neural tracks—motor neuron and cervical spinal cord—selected because these are the tissues in which *UNC13A* cryptic-exon inclusion has been observed [16]. SSU Δ resolve the two variants differently (Fig. 2c, left). The cryptic-exon variant (ES) increased usage at both the cryptic acceptor and donor (SSU Δ ≈ 2.3 × 10^−5^ and 5.2 × 10^−5^). The combined construct (B) produced a similar positive effect (3.3 × 10^−5^ and 7.6 × 10^−5^), whereas the matched exon control (EC) remained centered near zero (|SSU Δ| *<* 3 × 10^−8^). In contrast, the intronic risk variant (IS) produced a small negative change at both cryptic sites (≈ −6 × 10^−7^ and −2 × 10^−6^), while its matched intronic control (IC) again remained near zero (|SSU Δ| *<* 2 × 10^−8^). We further evaluated the tissue specificity of AG’s predictions. AG produced the same directional effects in cerebellar reference tracks (Fig. 2c, right); SSU Δ ≈ 1.4 × 10^−5^ and 3.1 × 10^−5^ for ES, comparable in magnitude to the neural tracks, even though cerebellum shows little cryptic-exon inclusion in the experimental data [6]. Smaller but still positive effects were additionally observed across the non-neural reference tracks, comprising T cell, lung fibroblast, umbilical-vein endothelial cell, and IMR-90 (Fig. 2c, right; ≈ 2.8 × 10^−6^ and 5.8 × 10^−6^ for ES). Consistent with this lack of tissue specificity, a broader track-level analysis showed that active predictions were distributed across diverse tissue and cell-type tracks (Appendix Fig. 6).

Because AG predictions can depend on the surrounding sequence supplied, we asked whether the predicted *UNC13A* effects were sensitive to context, repeating the analysis across windows from 2 kb to 1 Mb (Fig. 2d). Each variant retains its characteristic sign and approximate magnitude across the full range, with only minor, non-monotonic fluctuation and no systematic trend as more sequence is added. Placed against reference variant sets from Section 2.1, the *UNC13A* effects are not outliers (Table. 2e). The cryptic-exon variant reaches only the ~61st percentile of known Sqtl effect magnitudes and the intronic variant the ~6th, and both sit at similar percentiles within the matched non-sQTL control set.

Taken together, the *UNC13A* analysis supports partial rather than complete mechanistic fidelity. It recovers the qualitative signature. As the ES is introduced, the cryptic acceptor and donor gain both splice-site probability and predicted usage in the expected direction, while EC effects remain near zero. The IS is less consistent: in isolation it produces a small effect of the opposite sign at the cryptic sites, despite being a risk allele. The larger limitation is that the predicted effect lacks the context specificity of the established mechanism. Part of this shortfall, however, reflects the model’s intended scope. The *UNC13A* cryptic exon is de-repressed only upon TDP-43 loss of function—a trans-acting cellular state absent from the reference condition and never presented to the model. We regard this less as a failure of AG than as a boundary of the task as posed. Within this constraint, the model deserves credit for capturing the local splice architecture and part of the cis effect of the risk variants.

### 2.3 De novo inference: Regulatory hypothesis generation at ALS GWAS loci

We next move beyond retrospective validation to ask whether AG produces sufficiently specific regulatory signals to yield new biological insight into ALS. Unlike the previous settings, the goal here is not to recover a known target, but to ask whether AG can generate new biological hypotheses where the causal gene, regulatory element, and molecular effect are unknown. We analyzed seven ALS GWAS lead variants with candidate effector genes prioritized by spinal-cord QTL co-localization (Table 1) [17]; each locus is named following the original analysis in [17]. For each variant, the model scores a 1-Mb context at single-base resolution across multiple modalities. Each GWAS variant was compared with 25 matched local control variants and scored across three modalities—RNA-seq, CAGE, and DNase—each comprising multiple CNS-relevant biosample tracks grouped into spinal cord, frontal cortex, and other brain regions (e.g. brain, cerebellum and putamen). Out of 1-Mb window, a genomic position was called active when the GWAS variant’s predicted effect exceeded all of its matched controls and a control-derived magnitude threshold (Appendix B.3.3). The distribution of positional effect sizes illustrates the basis of this filtering strategy (Appendix Fig. 7a). The 95th and 99.9th percentile thresholds provide lenient and stringent filters for retaining positions with effects above the local background. Full details of locus selection, control matching, scoring, and transcript attribution are provided in Appendix B.3.

**Table 1.**
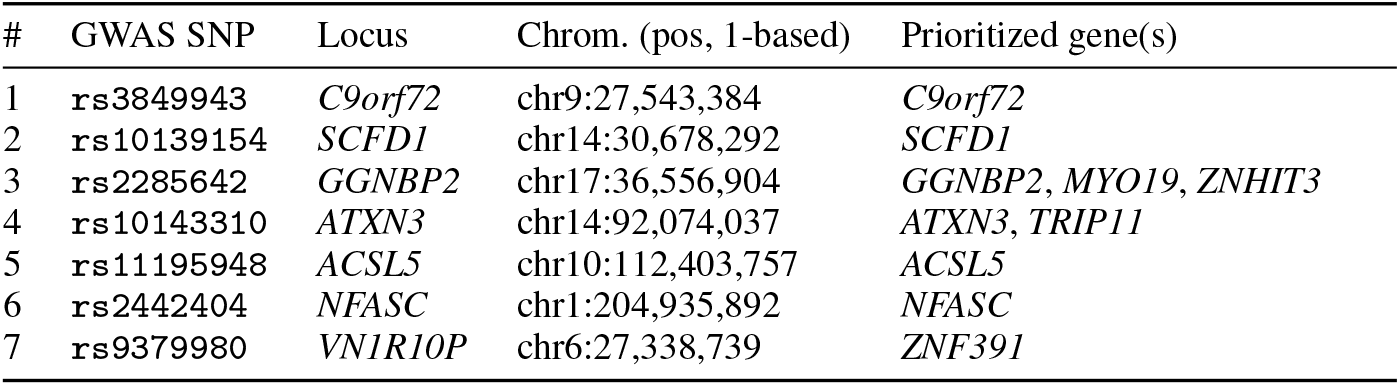
ALS GWAS loci used for the hypothesis generation analysis. Lead variants and previously prioritized candidate effector genes were obtained from spinal-cord QTL co-localization analysis reported by [17]. Following the original study, loci are named for the nearest protein-coding gene to the lead GWAS variant.

**Table 2.** Glossary of biological and genomics terminology used throughout the paper.

| Term | Definition |
| --- | --- |
| ALS | Amyotrophic lateral sclerosis; a fatal neurodegenerative disease causing progressive loss of motor neurons, leading to muscle weakness, paralysis, and eventually respiratory failure. |
| SNP / SNV | Single-nucleotide polymorphism / single-nucleotide variant; a change at a single position in the DNA sequence (e.g., an A swapped for a G). SNV is the general term for any such change; SNP typically refers to one common enough in the population to be called a "polymorphism." |
| Allele | One of the possible versions of a DNA sequence at a given position. A variant position typically has a *reference* allele (the standard/ancestral sequence) and an *alternate* allele (the variant form). |
| MAF | Minor allele frequency; how common the less-frequent allele of a variant is in a population (e.g., MAF = 0.05 means the rarer allele appears in 5% of chromosomes sampled). |
| GWAS | Genome-wide association study; a statistical approach that scans the genome for variants whose frequency correlates with a trait or disease across many individuals. |
| QTL | Quantitative trait locus; a genomic region (often tagged by a single variant) statistically associated with variation in a measurable trait, such as gene expression or splicing. |
| eQTL | Expression QTL; a variant associated with changes in how much a gene is expressed (transcribed). |
| sQTL | Splicing QTL; a variant associated with changes in how a gene's RNA transcript is spliced (i.e., which exons are joined together). |
| Splicing / splice site | The process by which non-coding regions (introns) are removed from RNA and coding regions (exons) are joined. A *splice site* is the specific sequence marking the boundary between an intron and exon; the *donor* site marks the start of an intron (5' end) and the *acceptor* site marks its end (3' end). |
| Splice junction | The connection point in mature RNA where two exons are joined after intron removal; different junctions correspond to different ways of splicing (isoforms). |
| Cryptic exon | An exon that is not normally included in the mature RNA transcript but becomes included when normal splicing regulation is disrupted (e.g., by loss of a regulatory protein). |
| Exon / Intron | Exons are the sequence segments retained in mature RNA (and typically encode protein); introns are segments removed during splicing. |
| TDP-43 | An RNA-binding protein that normally represses inclusion of certain cryptic exons; in ALS, TDP-43 mislocalizes out of the cell nucleus, causing loss of this repressive function and aberrant cryptic-exon inclusion (as seen in *UNC13A*). |
| *UNC13A* | A gene strongly associated with ALS risk; when TDP-43 function is lost, it develops a cryptic exon that disrupts normal protein production, making it a key model system for studying ALS splicing dysregulation. |
| *C9orf72*, *SCFD1*, *GGNBP2*, etc. | Names of specific genes located at ALS-associated genomic loci; each is a candidate "effector gene" thought to mediate disease risk at that locus. |
| Locus (loci) | A specific location/region in the genome, typically referring to a broader area around a variant or gene of interest (plural: loci). |

| **Term** | **Definition** |
| --- | --- |
| LD (Linkage disequilibrium) | The tendency of nearby genetic variants to be inherited together more often than expected by chance, making them statistically correlated even if only one is functionally causal. |
| Fine-mapping | Statistical methods used to narrow down a broad GWAS association signal to the most likely causal variant(s). |
| Heritability | The proportion of variation in a trait (e.g., ALS risk) within a population that can be attributed to genetic differences. |
| RNA-seq | RNA sequencing; a method for measuring which RNA transcripts are present and at what abundance in a sample. |
| CAGE | Cap analysis of gene expression; a sequencing method that identifies the precise transcription start sites of genes, used as a proxy for promoter/transcriptional activity. |
| DNase(-seq) | DNase I hypersensitive sites sequencing; a method that measures chromatin accessibility (how "open" DNA is), which correlates with regulatory activity. |
| Chromatin accessibility | How physically accessible a region of DNA is to regulatory proteins; open (accessible) chromatin is generally associated with active gene regulation. |
| Transcription factor | A protein that binds specific DNA sequences to help control (activate or repress) the transcription of nearby genes. |
| Promoter / TSS | A promoter is a DNA regulatory region where transcription is initiated; the TSS (transcription start site) is the exact position where RNA synthesis begins. |
| Biosample / Track | In genomic datasets, a "biosample" is a specific tissue or cell type profiled (e.g., spinal cord, motor neuron); a "track" is one specific dataset/measurement (e.g., one RNA-seq experiment) associated with that biosample. |
| Co-localization | A statistical method testing whether two association signals (e.g., a GWAS signal and an eQTL/sQTL signal) likely arise from the same underlying causal variant. |
| GRCh38 / hg38 | A standard reference version (build) of the human genome sequence, used to define consistent genomic coordinates. |
| 1000 Genomes Project | A public reference database of genetic variation from population sequencing, used here to determine allele frequencies and LD structure. |
| PLINK | A widely used software toolkit for analyzing genetic variation data, used here for LD pruning. |

We first asked whether ALS-associated regulatory activity localized to previously prioritized effector genes mentioned in [17], highlighted by green boxes in Fig. 3a. RNA local denotes activity in one exon in a transcript, whereas RNA body-wide denotes activity in at least two exons (Appendix B.3.4). At the *C9orf72* locus, AG provided little support for the prioritized gene itself. *C9orf72* itself carried almost no active signal; weak RNA-local activity in only two tissue contexts (25% of transcripts in frontal cortex, 38% in other brain regions) at the lenient threshold. The only consistent signal at the locus fell on the transcript *EMICERI* (50% of transcripts, RNA local), and this was the sole call to survive the stringent 99.9th-percentile threshold. This pattern held across other biosamples in Appendix Fig. 7b: *EMICERI* was active in RNA local across nearly every brain and spinal-cord track. The *SCFD1* locus showed the opposite pattern: here the signal fell on the prioritized gene. *SCFD1* was active across RNA, CAGE, and DNase outputs in both spinal cord and frontal cortex at the 95th-percentile threshold, and remained the associated gene under the stringent 99.9th-percentile threshold. This concentration on *SCFD1* held at individual-biosample resolution (Appendix Fig. 7b). The *GGNBP2* locus was more ambiguous because multiple genes at the locus had been nominated as candidate effectors in prior work [17]. At the lenient threshold, *GGNBP2* (up to 78–100% of transcripts) and *MYO19* (7–37%) were active together across tissues, with *ZNHIT3* contributing a weak signal (17%). Appendix Fig. 7b shows that this distributed signal persists; *GGNBP2, MYO19*, and *PIGW* were frequently active together across multiple biosamples and modalities.

**Figure 3.**
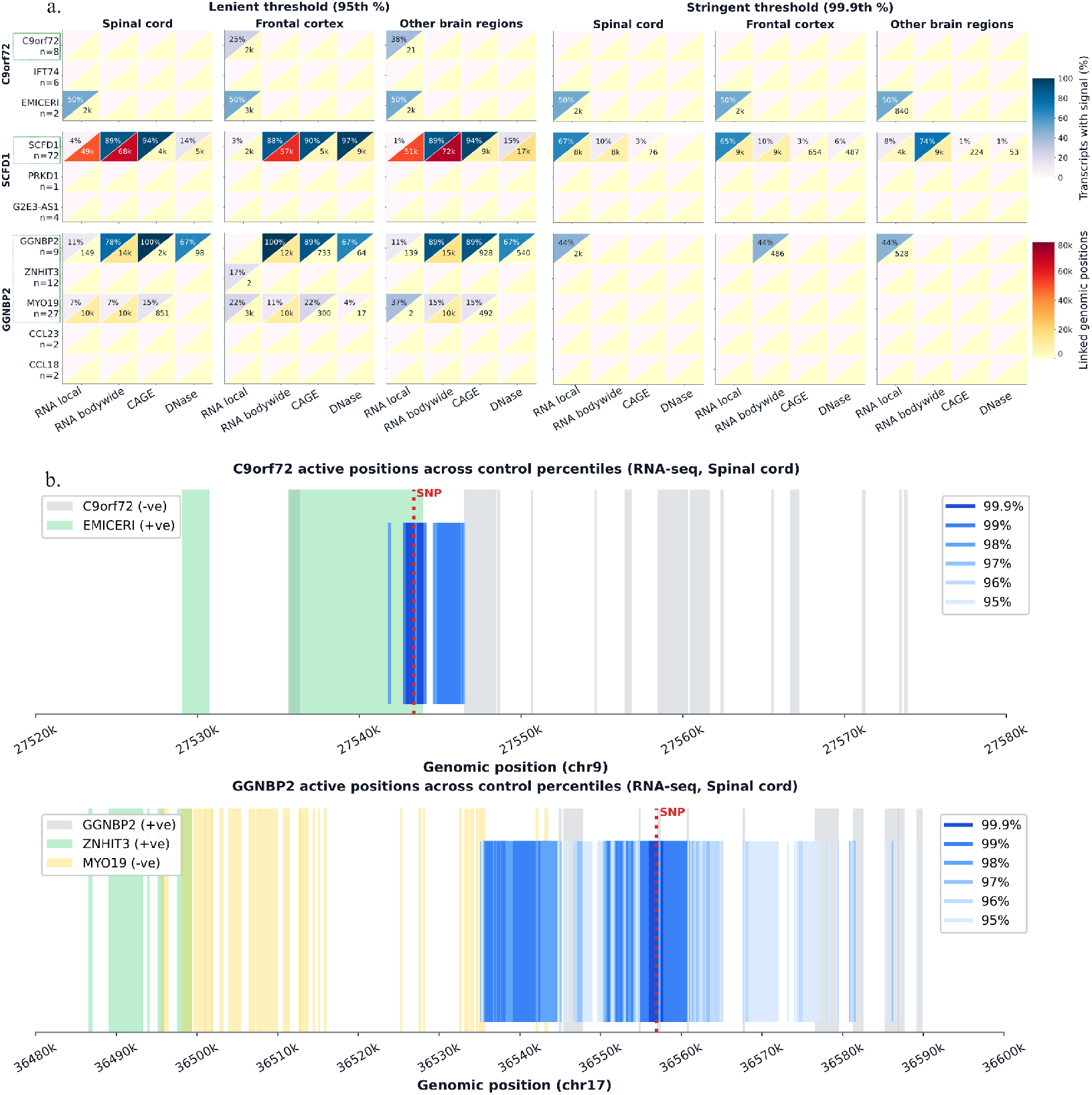
AG signal across control percentiles at ALS GWAS loci. AG signal at three ALS-associated loci (*C9orf72, SCFD1*, and *GGNBP2*). **(a)** Summary of active genomic positions at each locus. Rows are candidate genes (n = transcripts per gene). Columns group the AG output modalities (RNA local, RNA bodywide, CAGE, DNase), each shown across three tissue contexts (spinal cord, frontal cortex, other brain regions) and two activation thresholds (lenient, 95th percentile; stringent, 99.9th percentile of the control distribution). GWAS-nominated prioritized genes are green boxed. Each unit is split diagonally: the upper-left triangle (blue) gives the fraction of the gene’s transcripts with ≥ 1 active position, and the lower-right triangle (red) gives the number of linked genomic positions. **(b)** Active genomic positions along the *C9orf72* (chr9) and *GGNBP2* (chr17) loci (RNA-seq, spinal cord). Vertical blue bars mark active positions, shaded by the control effect percentile exceeded (95th–99.9th). Colored blocks mark the exons of each gene, keyed by color to the per-panel legend (± = strand); the red dashed line marks the ALS-associated SNP position.

The gene-level analysis revealed potentially novel signals. At the *C9orf72* locus, the neighboring transcript *EMICERI*, which was not among the prioritized effector genes, showed more consistent activity than *C9orf72* itself. Likewise, at the *GGNBP2* locus, the non-prioritized gene *PIGW* showed substantial AG signal (Appendix Fig. 7b). In both cases AG appeared to be pointing at genes that earlier prioritization had missed. We therefore examined whether these signals reflected gene-specific regulatory activity.

In both *C9orf72* and *GGNBP2*, the active RNA-seq positions were concentrated in a narrow region surrounding the GWAS SNP (Fig. 3b). At *C9orf72*, the active positions fell where *EMICERI* happens to overlap the SNP, while *C9orf72*’s own exons, several kilobases away, stayed silent. Across all seven loci, predicted effect size declines with a gene’s distance to the nearest exon (Fig. 4), and the prioritized gene scores highly only when it is the closest gene to the variant. At *GGNBP2* the gene ranking—*GGNBP2* > *MYO19* > *PIGW*—followed proximity to the variant rather than prioritization:

**Figure 4.**
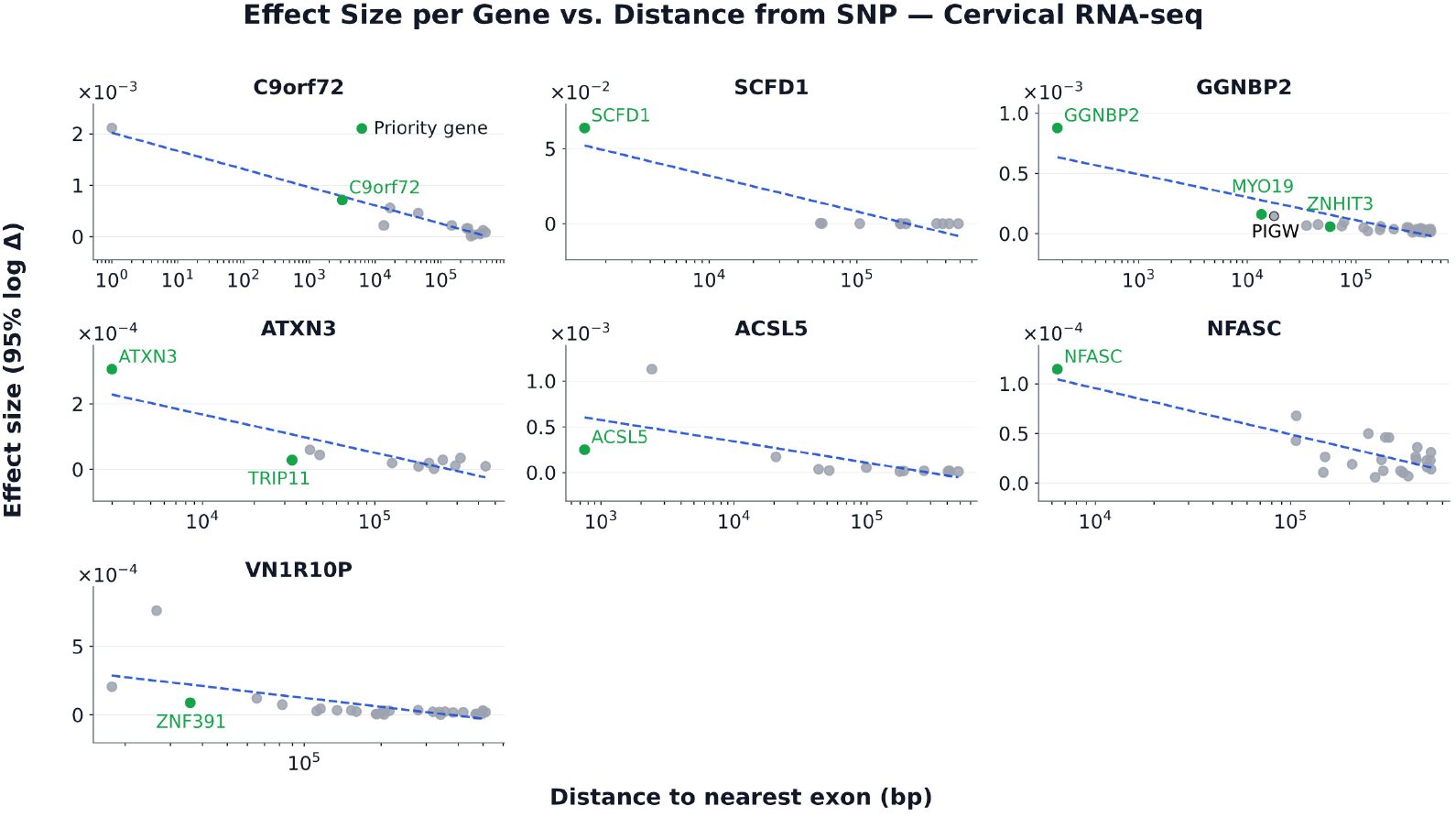
Predicted effect size versus distance to the nearest exon. Each panel plots the 95th-percentile predicted effect size (log Δ) for each gene against its distance from the GWAS SNP to the nearest exon. All protein-coding and lncRNA genes annotated within the 1-Mb scoring window are included: prioritized candidate genes in green, all other annotated genes in gray. Across all seven loci, predicted effect size declines with distance to the nearest exon (−0.95 SD per decade; pooled regression on effect sizes *z*-scored within each locus; *p* = 2.7 × 10^−23^).

**Figure 5.**
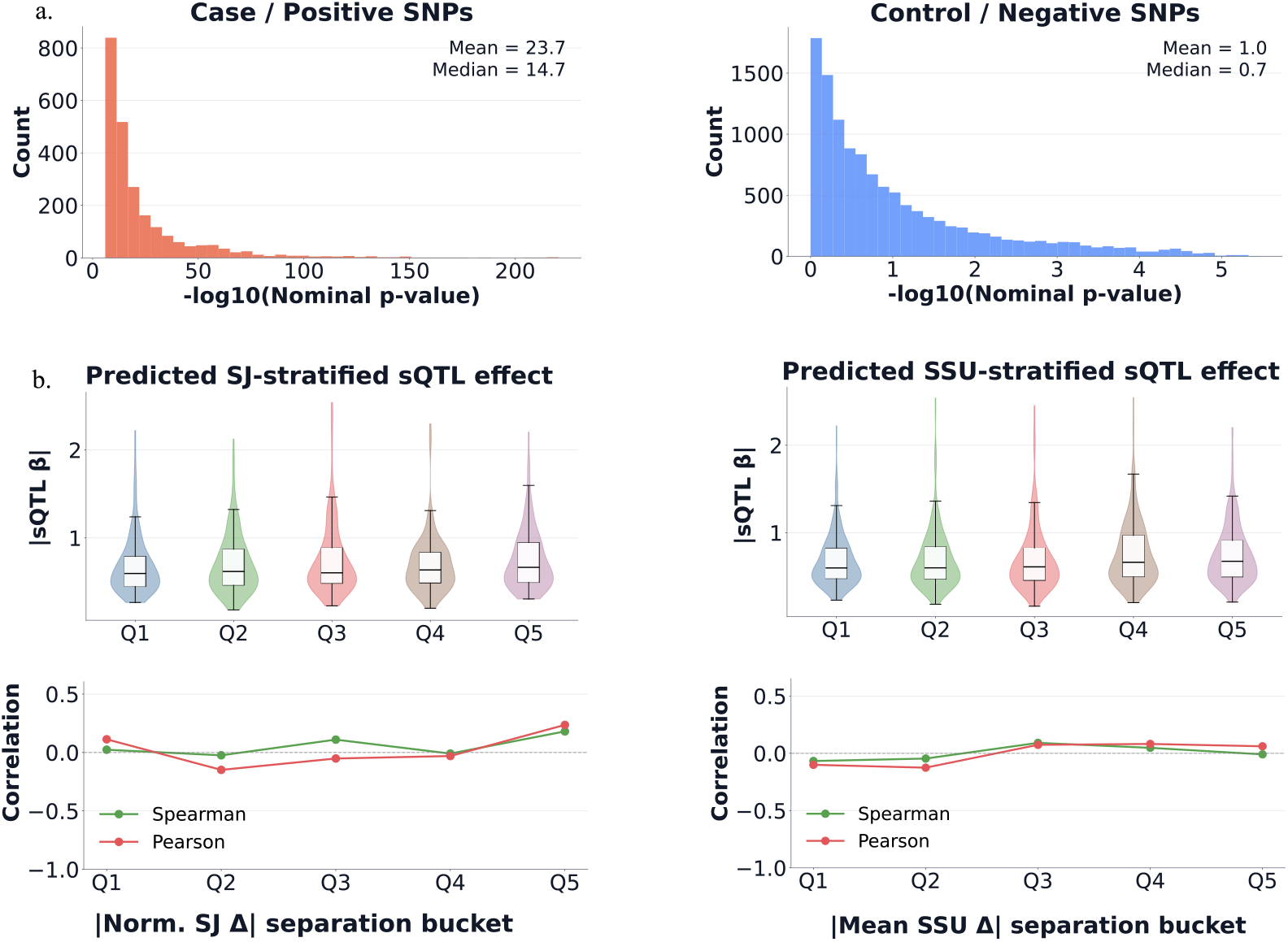
Relationship between predicted |Δ|_sep_ and measured sQTL. **(a)** Distribution of cervical sQTL significance (− log_10_ nominal *p*-value) from NYGC ALS controsium [16] for case/positive SNPs (left) versus control/negative SNPs (right); means and medians are annotated. The positive set is strongly significant (median 14.7) while matched controls are near-null (median 0.7), confirming the two groups are well separated on association strength. **(b)** Measured sQTL effect size (|*β*|) as a function of the predicted-effect separation (|Δ|_sep_, Eq. 2) bucket, for |Norm. SJ Δ| (left) and |Mean SSU Δ| (right). Top: distribution of |*β*| within each quintile bucket (Q1–Q5) of the predicted-effect separation statistic, shown as violins with overlaid box plots. Bottom: within-bucket Spearman (green) and Pearson (red) correlation between the AG-predicted effect and |*β*|. Correlations remain near zero across all buckets, indicating that predicted-effect magnitude does not track measured sQTL effect size.

**Figure 6.**
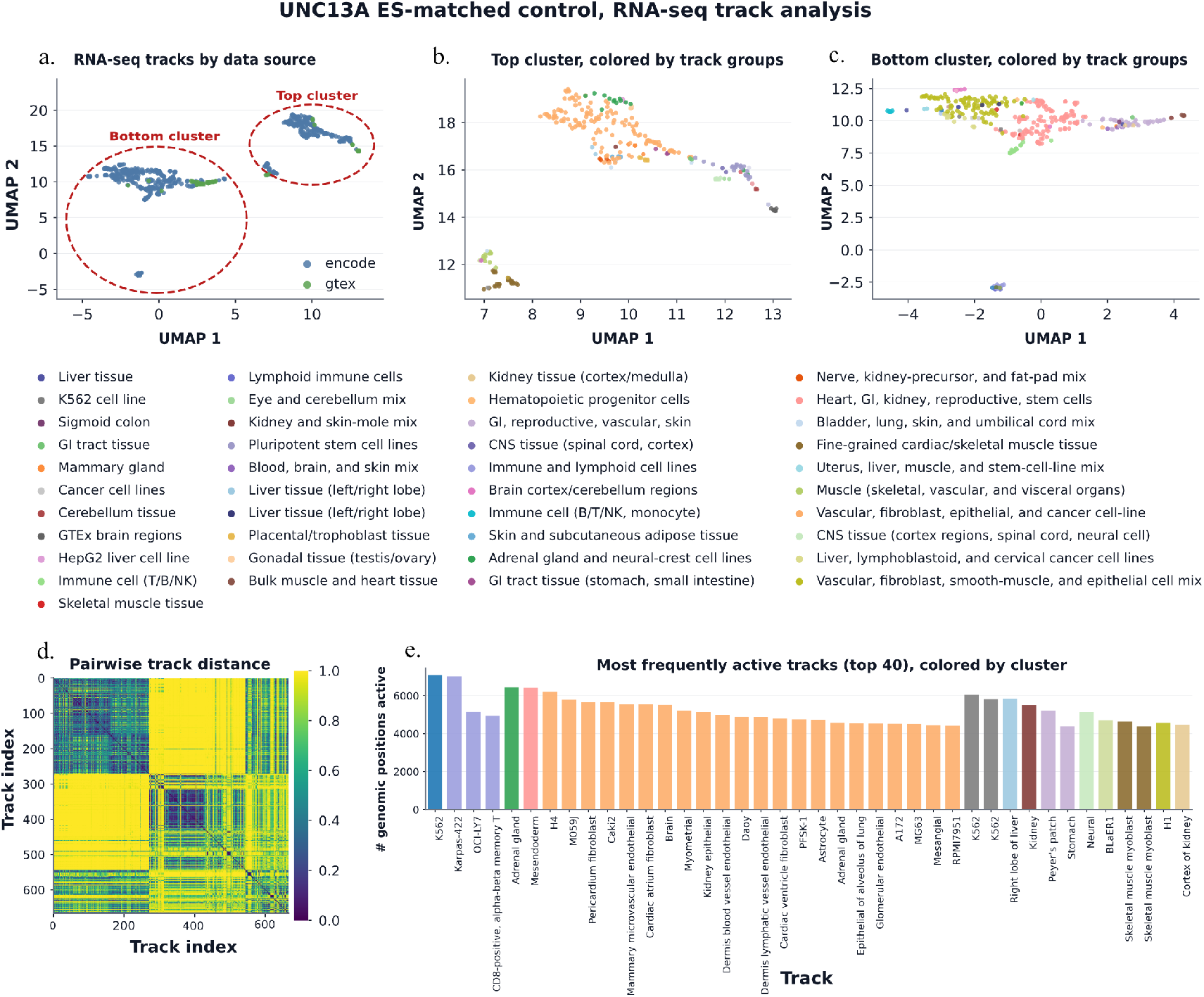
Control-derived organization of the UNC13A RNA-seq track space. Baseline organization of AG RNA-seq tracks derived entirely from the *UNC13A* exon risk variant (ES) matched-control (EC) analysis. To build this null geometry, we pooled all 25 EC controls across positions, correlated the RNA-seq tracks pairwise (Pearson on per-track *z*-scored effects), and defined a track distance *D* = 1 − |Corr|. Average-linkage hierarchical clustering and a UMAP embedding were then computed directly on *D*. Panels (a)–(d) describe this null, computed entirely from the EC with no ES activity involved; panel (e) overlays ES activity onto this fixed representation. **(a)** UMAP embedding of all RNA-seq tracks colored by data source (ENCODE or GTEx), revealing two broad control-defined clusters. **(b, c)** The top (b) and bottom (c) clusters re-embedded and colored by track group (shared legend). Fine-grained tissue groups do form local pockets of one color, so some tissue signal survives, but neither macro-cluster resolves into a coherent biological lineage: each spans many unrelated tissue and cell-type groups, indicating the null track geometry is broad and heterogeneous. **(d)** Pairwise distance matrix between RNA-seq tracks in EC, ordered to show the block structure underlying the control-derived clustering. **(e)** ES activity mapped onto the control-derived clusters. Each bar is one RNA-seq track; height is the number of positions called active for that track in the case-derived candidate mask (95th% control cutoff), and color is the track’s control-derived cluster. At this ALS-relevant locus, activity would be expected to preferentially occur in CNS, brain, and motor-neuron tracks. Instead, the most active tracks include immortalized cell lines (K562, Karpas-422, OCI-LY7), unrelated peripheral tissues (adrenal gland, kidney, muscle, liver) and span multiple control-derived clusters without neural enrichment. Even at the 95% threshold, AG recovers neither the expected tissue specificity nor coherent localization of the *UNC13A* cryptic-exon signal.

**Figure 7.**
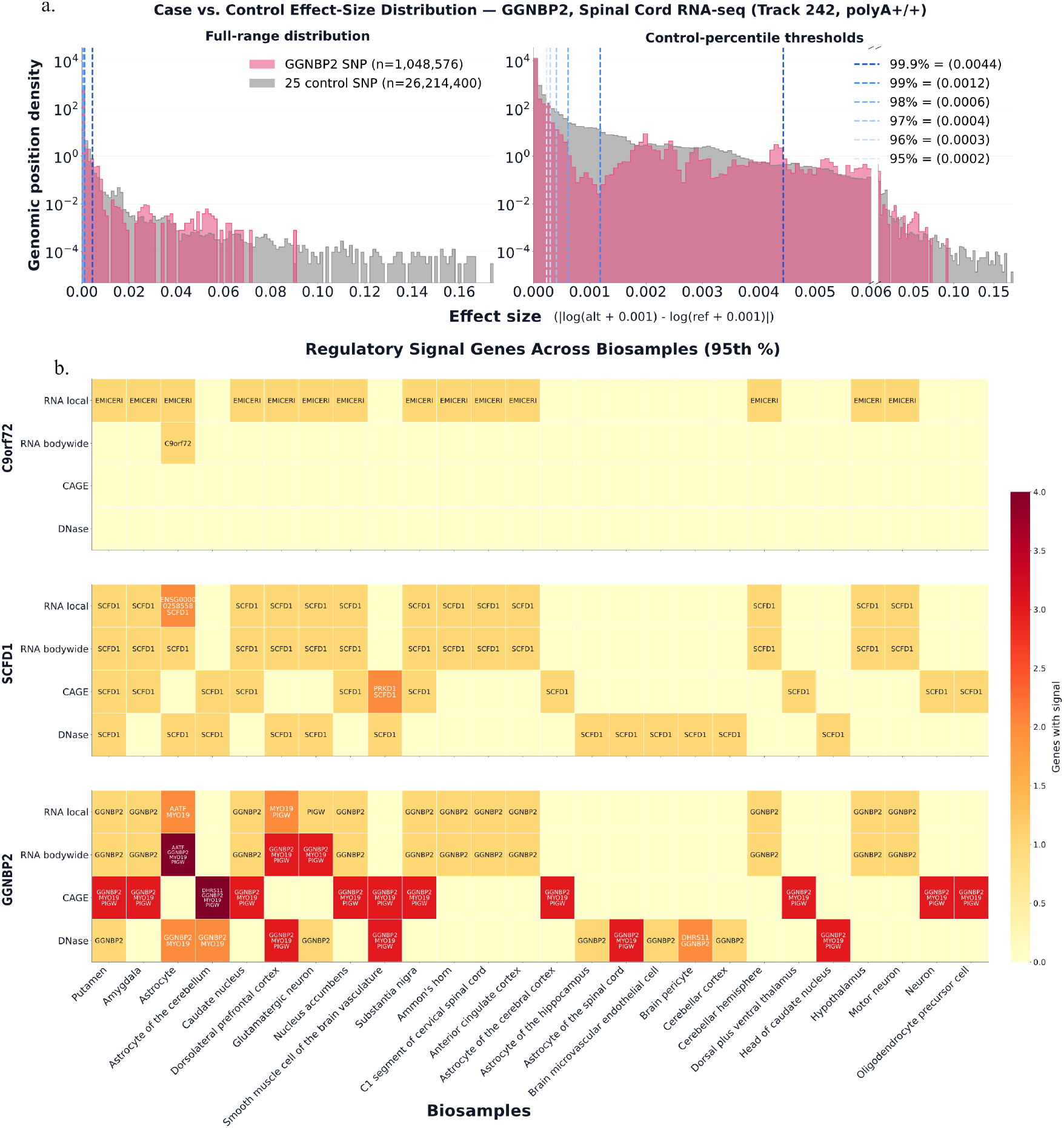
Case-versus-control AG predicted effect-size distributions and regulatory-signals across biosamples. **(a)** Distributions of the per-position AG predicted effect for the case variant (*GGNBP2* SNP, *n* = 1,048,576 genomic positions) versus its 25 pooled matched controls (*n* = 26,214,400), for a representative spinal-cord RNA-seq track (Track 242, polyA+), on full-range (left) and threshold-zoomed (right) axes. AG predicted effect size is | log(alt + 0.001) − log(ref + 0.001)|; dashed lines mark the control percentiles used as magnitude thresholds, with their absolute cutoffs (95th to 99.9th). Case and control distributions are largely superimposed, and the percentile cutoffs fall at small absolute effect sizes. **(b)** Genes called active at the 95th-percentile threshold across biosamples (columns) and AG modalities (rows) for the *C9orf72, SCFD1*, and *GGNBP2* loci; color encodes the number of genes active per unit. Active calls are sparse. They frequently fall on non-prioritized neighbors (*EMICERI* at *C9orf72*; *MYO19, PIGW, DHRS11* at *GGNBP2*) rather than the priority genes (Table. 1).

*PIGW* outranked *ZNHIT3* simply because it lay closer. This is a general pattern, not two isolated cases (Fig. 4). The “novel” signals at *EMICERI* and *PIGW* are thus not discoveries but proximity effects. Moreover each variant was scored over a 1,048,576,bp interval, yet at *GGNBP2* the fired positions span roughly 28,kb around the SNP, and at *C9orf72* only about 5,kb (Appendix Fig. 3b); in both, the active region occupies a few percent of the supplied megabase.

Of the three evaluations, de novo inference is the most consequential: it lacks ground truth, errors are hardest to detect, and validation requires costly wet-lab experiments. In this setting, the key failure is that AG signal largely reflects genomic proximity rather than specific regulatory biology. Predicted effects decay rapidly with distance, and active positions remain confined to a narrow neighborhood around the SNP despite the model receiving a 1-Mb sequence context. This makes gene-level interpretation particularly problematic for discovery.

## 3 Conclusion

AlphaGenome represents a remarkable advance toward general-purpose models of genome regulation and sets a strong benchmark for cis-regulatory prediction. It is already being used downstream to nominate candidate regulatory variants [25]. Our aim is not to discourage this direction, but to define the scope within which such predictions can currently support disease interpretation. On a zero-shot task it was never tuned for, AG flagged a subset of experimentally supported sQTLs with high confidence, localized both the canonical and cryptic splice sites at *UNC13A*, and predicted the direction of the exon-body risk variant correctly. Across three increasingly demanding evaluations, limitations emerge in accurately quantifying experimentally measured variant effects, recovering context-specific regulation, and distinguishing true effector genes from nearby alternatives. These results point to concrete opportunities for improvement: better calibration of variant-effect magnitude, stronger generalization to unseen variants, cell types, and genes, and greater discrimination of distal regulatory relationships from genomic proximity. Genomic foundation models are advancing rapidly; systematic disease-level evaluations such as this can help identify what additional capabilities are needed before their predictions can reliably support mechanistic discovery.

### Large Language Model Use

Large language models (LLMs) were used for language editing, clarification of technical concepts, and as a conversational tool for brainstorming and refining experimental ideas. All suggestions were critically evaluated by the authors.

## A Additional results

## B Method

### B.1 Quantitative fidelity: Data and Method

We constructed a sQTL benchmark to evaluate whether AlphaGenome (AG) predictions recover experimentally observed variant effects on RNA splicing in ALS-relevant human tissue. The analysis was organized into four stages: (i) construction of a positive sQTL set from the NYGC ALS Consortium sQTL resource [16]; (ii) construction of junction-specific matched negative variants; (iii) sequence-based scoring of each variant–junction pair; and (iv) evaluation of agreement with measured sQTL effects and discrimination of positive from negative variants.

#### B.1.1 Construction of the positive sQTL set

Positive/case variant–junction pairs were obtained from the NYGC ALS Consortium [16] sQTL: the primary region was cervical spinal cord (cervical_spinal). Significance was thresholded at *q <* 0.01 for the primary analysis. For each junction we kept only the lead variant. This choice prevented a single association signal from being represented repeatedly by multiple correlated variants in linkage disequilibrium (LD). The resulting positive label should therefore be interpreted as indicating that a variant tags a genetic signal associated with the splice junction; the selected variant is not necessarily itself the causal regulatory variant. Positives were filtered to single-nucleotide variants with minor allele frequency MAF ≥ 0.05 on the autosomes (chr1–22). A small number of variants were unscoreable because the fixed sequence window extended past a chromosome boundary. Starting from the lead-variant sQTL set, we applied three successive filters. First, matched non-sQTL controls were constructed for each junction; junctions for which no valid matched controls could be drawn were dropped. Second, each variant–junction pair was scored with AG on the fixed context window; pairs for which the variant and both splice sites could not be represented within this window were excluded. Third, junctions whose target splice junction could not be identified in AG’s splice-junction output were removed. After these steps, 1,254 positive SNP-junction groups remained for evaluation.

#### B.1.2 Construction of matched non-sQTL controls

Matched non-sQTL controls/negative variants let us ask whether AG assigns a larger effect to the experimentally supported sQTL than to comparable variants at the same junction that show no association, isolating regulatory signal from confounders such as splice-site proximity and allele frequency. Negative/control variants were drawn from nominal NYGC sQTL association results. The primary negative-set construction was performed for cervical spinal cord and restricted to junctions already represented in the positive set. For each positive junction, candidate negatives were variant–junction pairs whose nominal association *p*-value exceeded the junction-specific nominal significance threshold (Fig. 5a). Matching was performed independently within each junction. For each positive variant, candidate controls were matched using three features:

1. **1.Junction-span status** (exact match): whether the variant falls inside or outside the genomic span of the splice junction. A positive lying within the junction span was matched only to controls also lying within the span, and likewise for variants outside it, so that positives and controls share the same positional relationship to the junction.
2. **Variant–junction distance** (binned match): the distance from the variant to the splice junction, discretized into the bins [0, 50, 150, 400, 10^3^, 3 × 10^3^, 10^4^, 3 × 10^4^, 10^5^, ∞) bp. A control was eligible only if its distance fell in the same bin as the positive, ensuring that positives and controls are comparably close to the junction.
3. **Minor allele frequency** (binned match): the variant’s MAF in the reference panel, discretized into the bins [0.05, 0.10, 0.15, 0.25, 0.35, 0.50]. A control was eligible only if its MAF fell in the same bin as the positive, preventing systematic frequency differences between the two classes.

The final benchmark therefore used a strict 1:5 matched design: every retained positive sQTL had exactly five matched negative variants.

##### Linkage-disequilibrium filtering

To reduce the possibility that a nominally non-significant control tagged the same underlying association signal as its matched sQTL, LD pruning was performed pair-specifically within each junction with PLINK 2 [8]. Because NYGC whole-genome sequencing was inaccessible, the LD reference was the 1000 Genomes EUR panel (hg38; *n* = 525 samples [13]; 6,479,449 variants; MAF floor 0.05; *r*^2^ threshold 0.1). Panel joins were attempted under both variant-key orientations. Matching quality was checked with a Kolmogorov–Smirnov test on the distance and MAF distributions, requiring *p >* 0.05.

#### B.1.3 AG scoring and metric

Allele orientation was standardized before comparison with AG. The alternate allele was treated as the effect allele throughout the analysis, and AG effects were defined as

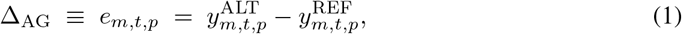

Where 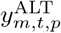 and 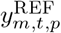 denote the AG prediction from the sequences carrying the alternate and reference alleles, respectively, for output modality *m*, biosample track *t*, and genomic position *p*. For classification analyses, we used the absolute predicted effect, |Δ_AG_|, because either an increase or decrease in predicted splicing can represent a regulatory perturbation. We additionally defined the |Δ|_sep_ within each matched junction group as the difference between the absolute AG effect assigned to the sQTL and the mean absolute effect assigned to its matched non-sQTL controls:

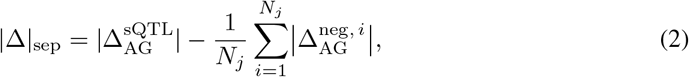

where 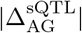 is the absolute AG effect (Eq. 1) assigned to the sQTL, 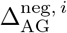 is that of its *i*-th matched control, and *N*_*j*_ is the number of matched controls for junction *j*. Thus, |Δ|_sep_ *>* 0 indicates that AG assigns the experimentally supported sQTL a larger predicted perturbation than the mean background effect among its matched controls, whereas |Δ|_sep_ *<* 0 indicates an effect below this junction-specific baseline.

Each variant was scored with AG on a fixed 131,072 bp window. Rather than centering the sequence window on the variant alone, we constructed a span containing the variant position together with the donor and acceptor coordinates of the associated junction. This ensured that the variant and both splice-junction endpoints were represented whenever compatible with the fixed sequence length and avoided unnecessary loss of the target junction for variants located asymmetrically relative to the splice sites. NYGC regions were mapped to AG biosamples under a fixed, reviewed correspondence: cervical spinal cord to the C1 cervical spinal cord segment (GTEx cervical spinal cord C1 poly(A)^+^ RNA-seq track UBERON:0006469; unstranded).

AG predicts several splicing-related outputs directly from sequence, and we derive three variant-effect scores from them. *(i) Normalized splice-junction score* (Norm. SJ Δ) is built from AG’s predicted splice-junction counts (`splice_junctions`). Because raw junction counts can vary with the overall predicted activity of the local splice region, we constructed a donor-normalized junction score. For a target junction *j* with donor *d*, let *C*_*j*_ denote its predicted junction count and let *A*(*d*) denote the set of acceptors competing with the same donor. The normalized junction score was defined as

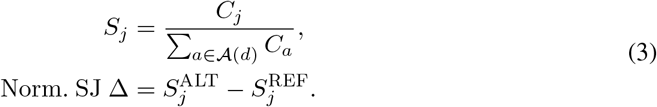

*(ii) Mean splice-site-usage score* Splice-site-usage (SSU) predictions provided a complementary endpoint. (Mean SSU Δ) is derived from AG’s predicted per-site usage (`splice_site_usage`); we average the usage change at the target junction’s donor and acceptor, giving a site-level measure of how strongly the variant perturbs the splice sites themselves. For each target splice site, the allele-specific change in predicted site usage was defined as

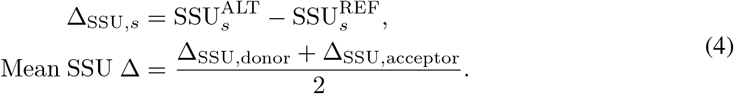

### B.2 Mechanistic fidelity: Data and Method

We used the well-characterized *UNC13A* cryptic-splicing mechanism as a locus-specific benchmark to test whether AG can recover a known ALS-associated regulatory perturbation. Unlike the genome-wide sQTL benchmark, where the lead associated variant may not itself be causal, the *UNC13A* experiment was designed around two established disease-associated variants and a predefined set of canonical and cryptic splice sites [20, 6]. We evaluated the variants individually, relative to matched local controls, across different sequence-context lengths, and jointly in a combined risk-allele sequence.

#### B.2.1 Construction of the positive variant and control

All analyses used GRCh38 coordinates for *UNC13A*, which lies on the negative strand of chromosome 19. Following [6], we curated the canonical exon 20/21 donor and acceptor and the cryptic-exon acceptor and donor bounding the CE interval. We evaluated two ALS-associated variants: intron variant (IS) rs12608932 at chr19:17,641,880, with protective allele A and risk allele C, and within the cryptic exon variant (ES) rs12973192 at chr19:17,642,430, with protective allele C and risk allele G.

##### Matched local controls

To determine whether the effects assigned to the two case variants were unusual relative to other variants in the same locus, we constructed separate sets of 25 matched control variants for IS and ES called intron control (IC) and exon control (EC), yielding 50 controls in total. Candidate controls were drawn from Esembl Variant Effect Predictor (VEP) [21] annotated set of healthy-control variants at the *UNC13A* locus and were required to carry no splice-related consequence, ensuring that controls did not themselves perturb the monitored splice sites. Matching, MAF filtering, and LD pruning against both case SNPs then followed the same framework used for the sQTL benchmark (Appendix B.1.2).

#### B.2.2 AG scoring and metric

We scored variants with AG on a 131,072 bp window and extracted predictions at the four curated splice sites (canonical donor and acceptor, cryptic acceptor and donor) from two splicing heads: splice-site probabilities (`splice_sites probs`) and splice-site-usage predictions (SSU Δ, `splice_site_usage`). *Splice-site probability* is AG’s predicted probability that a position functions as a splice site at all; we use it to ask whether the model recognizes a site as present, independent of how much it is used. Because *UNC13A* is minus-strand, only minus-strand tracks were used. To assess tissue specificity, effects were summarized over two ALS-relevant neural tracks — motor neuron and C1 cervical spinal cord — and contrasted against reference tracks not expected to support cryptic-exon inclusion: cerebellum, together with a non-neural set (T cell, lung fibroblast, umbilical-vein endothelial cell, and IMR-90). For each variant the alleles were oriented as protective versus risk, and the AG effect was defined as the paired difference under matched sequence context. To test whether the predicted *UNC13A* effect depended on the amount of surrounding genomic context provided to AG, we repeated the single-variant analysis for the two case variants across eight supported context sizes:

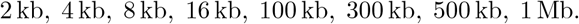

To assess whether the two variants act additively, we constructed a single combined perturbation carrying both risk alleles (B) against the matched both-protective sequence, scored on one shared window centered on the variant midpoint.

### B.3 De novo inference: Data and Method

The preceding benchmarks evaluate AG against known biological ground truth. Here, we instead test its prospective use at ALS GWAS loci with unresolved effector genes and mechanisms, asking whether control-calibrated AG signals are sufficiently specific to support new gene-level regulatory hypotheses.

#### B.3.1 Construction of the positive variant and control

We analyzed ALS GWAS loci and their candidate effector genes as reported by [17] (Table 1). Prioritization used co-localization of both eQTLs and sQTLs with the ALS GWAS signal in cervical and lumbar spinal cord. All coordinates are 1-based, GRCh38. Beyond these prioritized candidates, we scored every protein-coding and lncRNA gene annotated within the 1-Mb scoring window at each locus (GENCODE v46) [23], allowing AG to nominate regulatory targets independently of the prior QTL-based prioritization (Fig. 4). For each locus, 25 matched local controls were drawn from a VEP [21] annotated healthy-control variant set, excluding candidates with a splice-altering, protein-altering, or pathogenic consequence so that controls did not themselves carry a functional effect. Matching, MAF filtering, and LD pruning against the case variant then followed the same framework used for the sQTL benchmark (Appendix B.1.2).

#### B.3.2 AG scoring and metric

Each variant (case or control) was scored with AG on its locus’s shared 1,048,576 bp interval, restricted to three CNS-relevant modalities: RNA-seq (`rna_seq`), CAGE (`cage`), and DNase (`dnase`). Within each head we retained a reviewed subset of CNS/brain tracks (35 RNA-seq, 32 CAGE, and 17 DNase), spanning brain, cerebellum, spinal cord, frontal cortex, and related biosamples. For the coverage heads (`rna_seq, cage, dnase`), the per-track, per-position effect used the log-ratio of alternate to reference coverage,

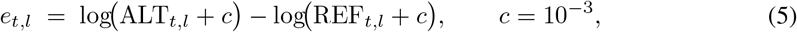

for track *t* and position *l*.

#### B.3.3 Empirical control-percentile calling of active positions

For each locus, let the ALS-associated GWAS SNP be the positive variant and let *K* = 25 denote its matched control variants, indexed by *k* = 1, …, *K*. AG predictions were evaluated across *M* = 3 modalities, *m* ∈ {RNA, CAGE, DNase}. Each modality contains *T*_*m*_ biosample tracks, indexed by *t*, and each prediction spans *L* = 1,048,576 genomic positions, indexed by *p* = 1, …, *L*.

Let

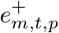

denote the AG allelic effect of the positive SNP at position *p* for modality *m* and track *t*, and let

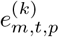

denote the corresponding effect of matched control *k* at the same genomic position. Because the positive SNP and all controls were evaluated over the same fixed locus interval, these effects could be compared position-by-position.

At each position, we first calculated the mean effect across the matched controls,

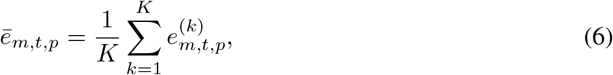

and the separation of the positive SNP from this local control background,

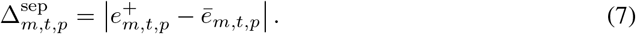

We also ranked the magnitude of the positive effect relative to the *K* matched controls at the same position,

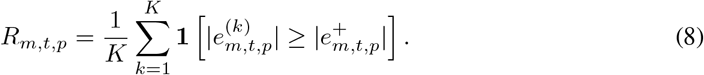

Thus, smaller *R*_*m,t,p*_ indicates that the positive SNP has a larger predicted effect than nearly all matched controls at that position.

Finally, for each modality–track pair we defined a magnitude threshold from the pooled matched-control predictions,

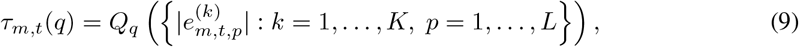

where *Q*_*q*_ denotes the *q*th percentile of the control distribution. A genomic position was called *active* when the positive SNP was more extreme than nearly all matched controls at that position and its separation from the control mean exceeded this empirical magnitude threshold:

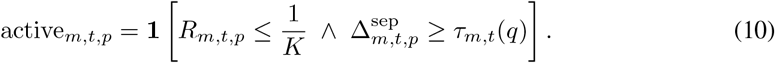

With *K* = 25, the rank criterion corresponds to *R*_*m,t,p*_ ≤ 0.04. We used *q* = 95 as the lenient threshold and *q* = 99.9 as the stringent threshold. For positional visualizations, we additionally evaluated *q* ∈ {95, 96, 97, 98, 99, 99.9}. Adjacent active 1-bp positions were merged into contiguous regions for downstream attribution and counting.

#### B.3.4 Transcript-level annotation and attribution

For each locus, we constructed a transcript-level annotation scaffold from GENCODE v46 [23], retaining protein-coding and lncRNA transcripts and their strand-aware transcription start sites (TSSs), promoters (TSS±1 kb), and exons. Active AG subregions were then attributed to transcripts according to the corresponding output modality. **RNA local** activity was assigned when an active RNA-seq subregion overlapped a single annotated exon of a transcript, whereas **RNA bodywide** activity required active RNA-seq signal across at least two distinct exons of the same transcript, indicating a broader transcript-level effect. **CAGE** activity was assigned when an active subregion fell within ±500 bp of the transcript TSS, and **DNase** activity when an active subregion overlapped the transcript promoter (TSS±1 kb). These rules provided a common transcript-level framework for relating modality-specific AG activity to candidate genes at each locus.

## References

[1] Žiga Avsec, Natasha Latysheva, Jun Cheng, Guido Novati, Kyle R Taylor, Tom Ward, Clare Bycroft, Lauren Nicolaisen, Eirini Arvaniti, Joshua Pan, et al. Advancing regulatory variant effect prediction with alphagenome. Nature, 649(8099):1206–1218, 2026.

[2] Chaima Ben Rabah and Ahmed Serag. Do foundation models truly outperform domain-specific models? evidence from digital pathology. Machine Learning and Knowledge Extraction, 8(6):164, 2026.

[3] Rishi Bommasani, Drew A Hudson, Ehsan Adeli, Russ Altman, Simran Arora, Sydney von Arx, Michael S Bernstein, Jeannette Bohg, Antoine Bosselut, Emma Brunskill, et al. On the opportunities and risks of foundation models. arXiv preprint arXiv:2108.07258, 2021.

[4] Danila Bredikhin, Alejandro Buendia, Martin Kjellberg, Christopher Zou, Xinming Tu, and Anshul Kundaje. Porting AlphaGenome to PyTorch. Genomics x AI Blog, March 2026. Published March 10, 2026.

[5] Garyk Brixi, Matthew G Durrant, Jerome Ku, Mohsen Naghipourfar, Michael Poli, Gwanggyu Sun, Greg Brockman, Daniel Chang, Alison Fanton, Gabriel A Gonzalez, et al. Genome modelling and design across all domains of life with evo 2. Nature, 652(8112):1349–1361, 2026.

[6] Anna-Leigh Brown, Oscar G Wilkins, Matthew J Keuss, Sarah E Kargbo-Hill, Matteo Zanovello, Weaverly Colleen Lee, Alexander Bampton, Flora CY Lee, Laura Masino, Yue A Qi, et al. Tdp-43 loss and als-risk snps drive mis-splicing and depletion of unc13a. Nature, 603(7899):131–137, 2022.

[7] Charlotte Bunne, Yusuf Roohani, Yanay Rosen, Ankit Gupta, Xikun Zhang, Marcel Roed, Theo Alexandrov, Mohammed AlQuraishi, Patricia Brennan, Daniel B Burkhardt, et al. How to build the virtual cell with artificial intelligence: Priorities and opportunities. Cell, 187(25):7045–7063, 2024.

[8] Christopher C Chang, Carson C Chow, Laurent CAM Tellier, Shashaank Vattikuti, Shaun M Purcell, and James J Lee. Second-generation plink: rising to the challenge of larger and richer datasets. Gigascience, 4(1):s13742–015, 2015.

[9] Johnathan Cooper-Knock, Sai Zhang, Kevin P Kenna, Tobias Moll, John P Franklin, Samantha Allen, Helia Ghahremani Nezhad, Alfredo Iacoangeli, Nancy Y Yacovzada, Chen Eitan, et al. Rare variant burden analysis within enhancers identifies cav1 as an als risk gene. Cell Reports, 33(9), 2020.

[10] Haotian Cui, Chloe Wang, Hassaan Maan, Kuan Pang, Fengning Luo, Nan Duan, and Bo Wang. scgpt: toward building a foundation model for single-cell multi-omics using generative ai. Nature methods, 21(8):1470–1480, 2024.

[11] Tong Ding, Sophia J Wagner, Andrew H Song, Richard J Chen, Ming Y Lu, Andrew Zhang, Anurag J Vaidya, Guillaume Jaume, Muhammad Shaban, Ahrong Kim, et al. A multimodal whole-slide foundation model for pathology. Nature medicine, 31(11):3749–3761, 2025.

[12] Michael Eisenstein. ‘virtual cells’ aim to turn raw data into predictive models of biology. Nature, 654(8117):286–288, 2026.

[13] Susan Fairley, Ernesto Lowy-Gallego, Emily Perry, and Paul Flicek. The international genome sample resource (igsr) collection of open human genomic variation resources. Nucleic acids research, 48(D1):D941–D947, 2020.

[14] Judit García-González and Krzysztof Gogolewski. Alphagenome: a swiss-army knife for exploring non-coding dna. Trends in Genetics, 2025.

[15] Orla Hardiman, Ammar Al-Chalabi, Adriano Chio, Emma M Corr, Giancarlo Logroscino, Wim Robberecht, Pamela J Shaw, Zachary Simmons, and Leonard H Van Den Berg. Amyotrophic lateral sclerosis. Nature reviews Disease primers, 3(1):17071, 2017.

[16] Jack Humphrey, Ali Oku, Marta Byrska-Bishop, Anna O Basile, Uday S Evani, André Corvelo Alex Tokolyi, Kailash Bp, Aline Réal, Yebin Kim, et al. The new york genome center als consortium resource integrates postmortem tissue transcriptomics and whole genome sequencing to empower biological discovery. medRxiv, pages 2026–04, 2026.

[17] Jack Humphrey, Sanan Venkatesh, Rahat Hasan, Jake T Herb, Katia de Paiva Lopes, Fahri Küçükali, Marta Byrska-Bishop, Uday S Evani, Giuseppe Narzisi, Delphine Fagegaltier, et al. Integrative transcriptomic analysis of the amyotrophic lateral sclerosis spinal cord implicates glial activation and suggests new risk genes. Nature Neuroscience, 26(1):150–162, 2023.

[18] Alfredo Iacoangeli, Isabella Fogh, Sashika Selvackadunco, Simon D Topp, Aleksey Shatunov, Wouter van Rheenen, Ahmad Al-Khleifat, Sarah Opie-Martin, Antonia Ratti, Andrea Calvo, et al. Scfd1 expression quantitative trait loci in amyotrophic lateral sclerosis are differentially expressed. Brain communications, 3(4):fcab236, 2021.

[19] Zeming Lin, Halil Akin, Roshan Rao, Brian Hie, Zhongkai Zhu, Wenting Lu, Nikita Smetanin, Robert Verkuil, Ori Kabeli, Yaniv Shmueli, et al. Evolutionary-scale prediction of atomic-level protein structure with a language model. Science, 379(6637):1123–1130, 2023.

[20] X Rosa Ma, Mercedes Prudencio, Yuka Koike, Sarat C Vatsavayai, Garam Kim, Fred Harbinski, Adam Briner, Caitlin M Rodriguez, Caiwei Guo, Tetsuya Akiyama, et al. Tdp-43 represses cryptic exon inclusion in the ftd–als gene unc13a. Nature, 603(7899):124–130, 2022.

[21] William McLaren, Laurent Gil, Sarah E Hunt, Harpreet Singh Riat, Graham RS Ritchie, Anja Thormann, Paul Flicek, and Fiona Cunningham. The ensembl variant effect predictor. Genome biology, 17(1):122, 2016.

[22] Michael Moor, Oishi Banerjee, Zahra Shakeri Hossein Abad, Harlan M Krumholz, Jure Leskovec, Eric J Topol, and Pranav Rajpurkar. Foundation models for generalist medical artificial intelligence. Nature, 616(7956):259–265, 2023.

[23] Jonathan M Mudge, Sílvia Carbonell-Sala, Mark Diekhans, Jose Gonzalez Martinez, Toby Hunt, Irwin Jungreis, Jane E Loveland, Carme Arnan, If Barnes, Ruth Bennett, et al. Gencode 2025: reference gene annotation for human and mouse. Nucleic acids research, 53(D1):D966–D975, 2025.

[24] Marie Ryan, Mark Heverin, Russell L McLaughlin, and Orla Hardiman. Lifetime risk and heritability of amyotrophic lateral sclerosis. JAMA neurology, 76(11):1367–1374, 2019.

[25] Ralf Tambets, Mihkel Jesse, Jaanika Kronberg, Adriaan van der Graaf, Erik Abner, Urmo Võsa, Ida Rahu, Nele Taba, Anastassia Kolde, Dzvenymyra Yarish, et al. Genetic analysis of circulating metabolic traits in 619,372 individuals. Nature, pages 1–8, 2026.

[26] Betina B Trabjerg, Fleur C Garton, Wouter van Rheenen, Fang Fang, Robert D Henderson, Preben Bo Mortensen, Esben Agerbo, and Naomi R Wray. Als in danish registries: Heritability and links to psychiatric and cardiovascular disorders. Neurology: Genetics, 6(2):e398, 2020.

[27] Wouter Van Rheenen, Aleksey Shatunov, Annelot M Dekker, Russell L McLaughlin, Frank P Diekstra, Sara L Pulit, Rick AA Van Der Spek, Urmo Võsa, Simone De Jong, Matthew R Robinson, et al. Genome-wide association analyses identify new risk variants and the genetic architecture of amyotrophic lateral sclerosis. Nature genetics, 48(9):1043–1048, 2016.

[28] Wouter Van Rheenen, Rick AA Van Der Spek, Mark K Bakker, Joke JFA Van Vugt, Paul J Hop, Ramona AJ Zwamborn, Niek De Klein, Harm-Jan Westra, Olivier B Bakker, Patrick Deelen, et al. Common and rare variant association analyses in amyotrophic lateral sclerosis identify 15 risk loci with distinct genetic architectures and neuron-specific biology. Nature genetics, 53(12):1636–1648, 2021.

[29] Eugene Vorontsov, Alican Bozkurt, Adam Casson, George Shaikovski, Michal Zelechowski, Kristen Severson, Eric Zimmermann, James Hall, Neil Tenenholtz, Nicolo Fusi, et al. A foundation model for clinical-grade computational pathology and rare cancers detection. Nature medicine, 30(10):2924–2935, 2024.

[30] Hanchen Wang, Tianfan Fu, Yuanqi Du, Wenhao Gao, Kexin Huang, Ziming Liu, Payal Chandak, Shengchao Liu, Peter Van Katwyk, Andreea Deac, et al. Scientific discovery in the age of artificial intelligence. Nature, 620(7972):47–60, 2023.

[31] Lily Wang, Peilin Jia, Russell D Wolfinger, Xi Chen, and Zhongming Zhao. Gene set analysis of genome-wide association studies: methodological issues and perspectives. Genomics, 98(1):1–8, 2011.

[32] Ali Yousefian-Jazi, Min Kyung Sung, Taeyeop Lee, Yoon-Ho Hong, Jung Kyoon Choi, and Jinwook Choi. Functional fine-mapping of noncoding risk variants in amyotrophic lateral sclerosis utilizing convolutional neural network. Scientific Reports, 10(1):12872, 2020.

[33] Sai Zhang, Johnathan Cooper-Knock, Annika K Weimer, Minyi Shi, Tobias Moll, Jack NG Marshall, Calum Harvey, Helia Ghahremani Nezhad, John Franklin, Cleide dos Santos Souza, et al. Genome-wide identification of the genetic basis of amyotrophic lateral sclerosis. Neuron, 110(6):992–1008, 2022.

